# Design, immunogenicity and efficacy of a Pan-SARS-CoV-2 synthetic DNA vaccine

**DOI:** 10.1101/2021.05.11.443592

**Authors:** Charles C. Reed, Katherine Schultheis, Viviane M. Andrade, Richa Kalia, Jared Tur, Blake Schouest, Dustin Elwood, Jewell N. Walters, Igor Maricic, Arthur Doan, Miguel Vazquez, Zeena Eblimit, Patrick Pezzoli, Dinah Amante, Maciel Porto, Brandon Narvaez, Megan Lok, Brittany Spence, Heath Bradette, Heather Horn, Maria Yang, Joseph Fader, Roi Ferrer, David B. Weiner, Swagata Kar, J. Joseph Kim, Laurent M. Humeau, Stephanie J. Ramos, Trevor R.F. Smith, Kate E. Broderick

## Abstract

Here we have employed SynCon^®^ design technology to construct a DNA vaccine expressing a pan-Spike immunogen (INO-4802) to induce broad immunity across SARS-CoV-2 variants of concern (VOC). Compared to WT and VOC-matched vaccines which showed reduced cross-neutralizing activity, INO-4802 induced potent neutralizing antibodies and T cell responses against WT as well as B.1.1.7, P.1, and B.1.351 VOCs in a murine model. In addition, a hamster challenge model demonstrated that INO-4802 conferred superior protection following intranasal B.1.351 challenge. Protection against weight loss associated with WT, B.1.1.7, P.1 and B.1.617.2 challenge was also demonstrated. Vaccinated hamsters showed enhanced humoral responses against VOC in a heterologous WT vaccine prime and INO-4802 boost setting. These results demonstrate the potential of the pan-SARS-CoV-2 vaccine, INO-4802 to induce cross-reactive immune responses against emerging VOC as either a standalone vaccine, or as a potential boost for individuals previously immunized with WT-matched vaccines.

## Introduction

COVID-19 remains a global pandemic. To date SARS-CoV-2 has infected over 185 million people, and over 4 million people have succumbed to disease [1]. Concerningly, SARS-CoV-2 variants containing novel mutations impacting virological and epidemiological characteristics are driving an increased level COVID-19 morbidity and mortality in many parts of the world [2-4]. Several variants have become a focus of attention. The B.1.1.7 (Alpha), B.1.351 (Beta), P.1 (Gamma) and B.1.617.2 (Delta) variants have rapidly become dominant in some regions (reviewed in [5]). Importantly, some of the mutations associated with these VOCs enhance resistance to neutralizing antibodies induced after infection or vaccination [6-11]. Recent clinical data has revealed a significant decrease in efficacy of vaccines against novel mutations in the SARS-CoV-2 spike protein [6, 7, 12]. A recent Phase 3 clinical trial investigating the Vaxzevria vaccine demonstrated low efficacy (21.9%) against the circulating B.1.351 VOC in South Africa [13]. Additionally, reports of fully vaccinated individuals suffering breakthrough infections with SARS-CoV-2 variants further highlight the risk variants pose [14, 15]. Pseudotyped virus and live virus neutralization results have showed reduced or in some cases loss of neutralizing antibody capacity against the B.1.351 and B.1.617.2 variants. Multiple growing lines of evidence are converging on the need to effectively address new variants and for strategies to refine vaccine designs for further emerging VOC challenges [16, 17].

Vaccine strategies to address the critical global challenge of VOCs are being developed and tested. These include immunogens based on an individual variant (matched strain) or designs that aim to provide pan-SARS-CoV-2 variant coverage [18, 19]. We utilized our SynCon^®^ DNA-based vaccine technology for rapid design, generation, and synthesis of new vaccine candidates to meet the challenge of emerging SARS-CoV-2 variants. The vaccine candidates were generated based on surveys of emerging variants with the goal of generating a pan-SARS-CoV-2 antigen which can demonstrate effective cross protection to multiple variants of concern. From the candidate pool we selected a vaccine which encodes a novel set of SARS-CoV-2 spike changes reflective of globally observed mutations from multiple lineages and containing critical receptor binding domain (RBD) changes present in variants of concern circulating worldwide. In previous studies, immunization of small and large animal models with DNA vaccines encoding SARS-CoV-2 spike protein (INO-4800, currently in Phase 2/3 clinical trials) provided protection against disease challenge with the matched virus [20-22].

Here, we report on the design, broad immunogenicity and efficacy of a next-generation pan-SARS-CoV-2 vaccine, INO-4802 and its use in prime and heterologous boost regimens targeting emerging SARS-CoV-2 variants of concern in preclinical animal models. Studies highlight the use of rational design processes to identify a single immunogen encoded in plasmid DNA and delivered by CELLECTRA® electroporation which can induce broadly cross-reactive neutralizing antibodies against SARS-CoV-2 variants.

## Results

### Design strategy of a pan-SARS-CoV-2 vaccine

The SynCon design strategy employed to create a pan-SARS-CoV-2 vaccine candidate is described in a step-by-step manner (**Fig. 1a**). Our design goal was to drive neutralizing coverage against multiple VOCs by providing maximal overlap to critical regions of multiple current VOCs. SARS-CoV-2 genome sequence entries (derived from GISAID) covering a four-month period (October 2020-January 2021) were sampled from multiple geographic regions (Brazil, Canada, India, Italy, Japan, Nigeria, South Africa, United Kingdom, United States) to provide a broadly representative pool of current and emerging variants. Consistent mutations in the SARS-CoV-2 Spike sequences were aggregated for each region. The survey detected large numbers of low frequency mutations which if included wholesale would result in a sequence too divergent from real circulating variants. To prevent this, we manually curated and selected mutations for inclusion. Any low frequency mutation was only considered if it was widespread across multiple geographical locations. The results from each of these regions were then aggregated to determine a common set of overlapping mutations. The number of mutations selected was informed from past experience in generating SynCon sequences as well as knowledge of SARS-CoV-2 spike regions which contribute the majority of neutralizing epitopes, in particular the N-terminal domain (NTD) and receptor binding domain (RBD) [23-25]. Manual sequence inspection and observations derived from spike molecular models informed decisions on number and placement of mutations from the pool of aggregated mutations (**Fig. 1b**). We avoided making large numbers of changes, keeping spike sequence identity at about 99% based on cross-comparisons of WT and VOC spike sequences. By design we avoided aggregating large numbers of mutations that did not naturally co-occur to reduce the potential of generating novel non-relevant epitopes. For ease of understanding, all mutations and changes are numbered according to the canonical SARS-CoV-2 spike sequence numbering scheme. Additional amino acid changes were added in the receptor binding domain (RBD) to shift the RBD to be more like VOCs that are currently known to escape neutralizing antibodies [6-10]. A tandem proline mutation (K986P/V987P) named “2P” was added to the SynCon SARS-CoV-2 Spike to augment protein stability [26, 27]. Sequence changes were mapped schematically as well as onto a spike trimer model containing the changes (**Fig. 1a** and **b**). The final single construct containing all changes was termed INO-4802. The design strategy results can be visualized using an unrooted phylogenetic tree comparing the spike sequences of INO-4802 to several other constructs used in the studies along with several circulating lineages including multiple VOCs (**Fig. 1c**). INO-4802 clusters closely but does not overlap with any VOCs. Plasmids matched to WT (pWT) and B.1.351 (pB.1.351) variants were used as control immunogens in this study, they show identity to the matched Spike glycoprotein sequences as expected (**Table 1**). GISAID accession numbers of entries used in the analysis are available in the Supplemental Materials.

**Figure 1.**
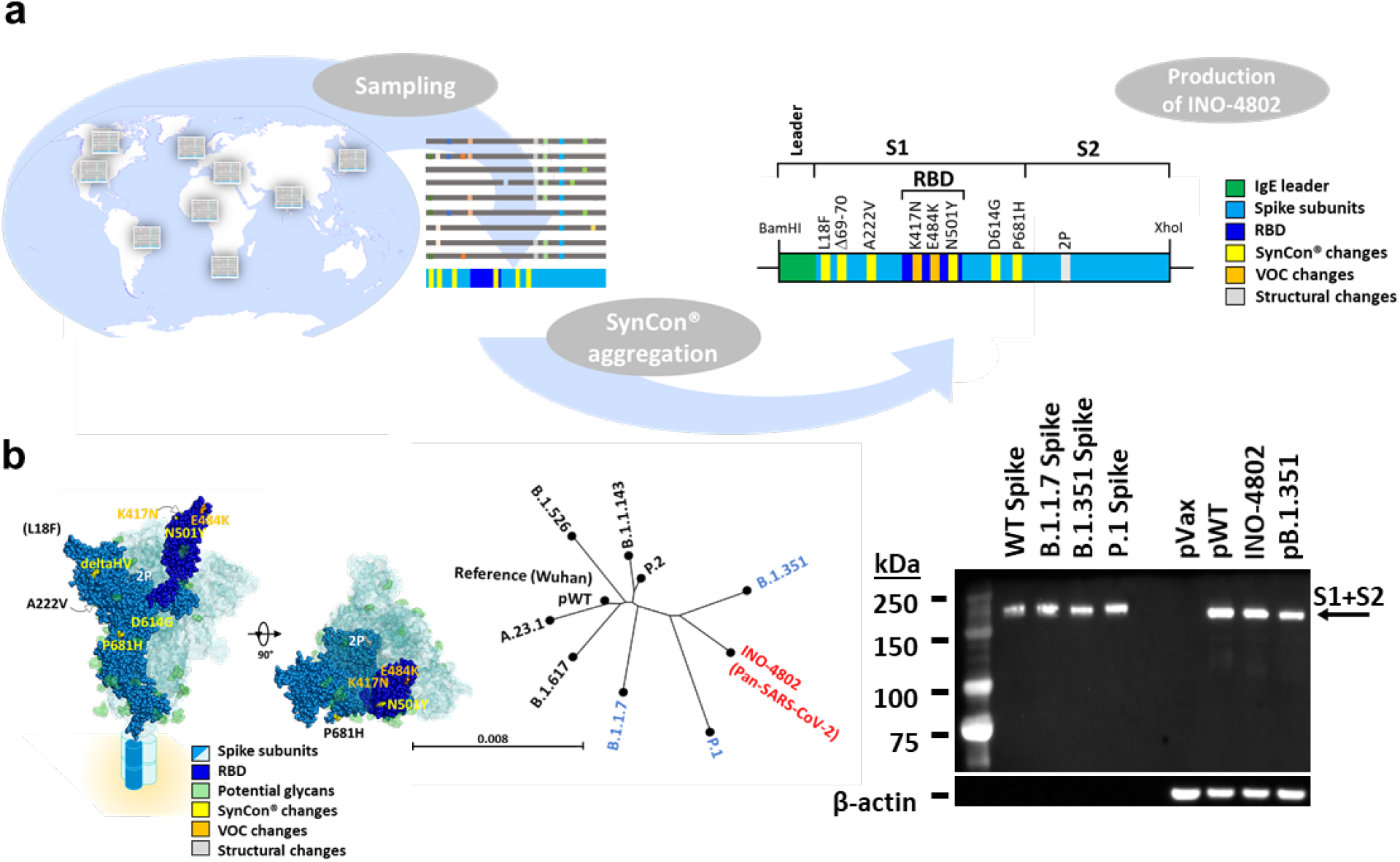
Design strategy of a pan-SARS-CoV-2 vaccine. **a)** SARS-CoV-2 spike glycoprotein sequences were sampled from multiple countries (Brazil, Canada, India, Italy, Japan, Nigeria, South Africa, the United Kingdom, and the United States) and the most prevalent mutations were aggregated for each location. The regional mutations were analyzed and aggregated to generate the SynCon spike. Lastly, in addition to the global SynCon changes to the spike (yellow), supplemental variant of concern (VOC) changes (orange) *was* placed in the receptor binding domain (RBD, dark blue) and the 2P mutation was added (indicated in gray) producing INO-4802 (pan-SARS-CoV-2). **b)** Molecular model of SARS-CoV-2 spike showing locations of mutations for INO-4802 colored similarly to 1a. For clarity a single spike subunit is labeled. Remaining subunits are indicated as transparent surfaces. Mutations not easily visible on a view are indicated with arrows and not all mutations are indicated in all views. Orthogonal views of the model are shown. Potential glycosylation sites are indicated in light green. The L18F mutation is not visualized in the model. The stalk region and membrane orientation are indicated by cartoon schematic. **c)** An unrooted phylogenetic tree comparing protein sequences derived from INO-4802, pB.1.351, and pWT spikes (black, gray, and red respectively) as well as spike sequences from a sampling of circulating variants (black) including current VOCs (blue). **d)** Analysis of in vitro expression of Spike protein after transfection of 293T cells with empty vector (pVax), pWT, INO-4802, or pB.1.351 plasmid by Western blot. Control proteins and 293T cell lysates were resolved on a gel and probed with a polyclonal anti-SARS-CoV-2 Spike RBD Protein. Blots were stripped then probed with an anti-β-actin loading control. Bands were detected at the expected SARS-COV-2 Spike antigen molecular weight of about 180 kDa inclusive of glycosylation.

**Table 1.** Vaccine construct descriptions.

| Vaccine Construct | Mutations | Design |
| --- | --- | --- |
| pWT | Original Wuhan sequence (NC_045512) | Matched to Wuhan sequence |
| pB.1.351 | L18F/D80A/D215G/Δ242-244/R246I/K417N/E484K/N501Y/D614G/A701V | Matched to B.1.351 |
| INO-4802 | L18F/Δ69-70/A222V/K417N/E484K/N501Y/D614G/P681H/K986P/V987P | Pan-SARS-CoV-2 |
| pVax | - | Empty insert plasmid control |

### *In vitro* expression of immunogens

To confirm expression of the immunogens we measured *in vitro* Spike protein production in HEK-293T cells after transfection with the corresponding plasmid constructs by Western blot analysis using a cross-reactive antibody against the RBD region of the SARS-CoV-2 S protein on cell lysates. Western blots of the lysates of HEK-293T cells transfected with pWT, pB.1.351, or INO-4802 constructs revealed bands approximate to the Spike protein molecular weight (**Fig. 1d**). Similar results were observed when using antibodies specific to the S1 or S2 regions of the spike as well (**Supplemental Fig. 1a**). Spike transgene RNA expression was also confirmed by RT-PCR analysis of COS-7 cells transfected with pWT, pB.1.351, and INO-4802 plasmids (**Supplemental Fig. 1b**). In summary, *in vitro* studies revealed the expression of the Spike transgene at both the RNA and protein level after transfection of cell lines with all three candidate vaccine constructs.

### Vaccination with INO-4802 induces binding and neutralizing antibodies against SARS-CoV-2 variants

The immunogenicity of the pWT, pB.1.351, and INO-4802 DNA vaccines was evaluated in the BALB/c mouse model. Mice were dosed with 10 µg plasmid DNA (pDNA) on day 0 and 14, and sera samples were collected on day 21 for evaluation. IgG binding titers against the full Spike protein of the WT and variants including B.1.1.7, P.1, and B.1.351 were evaluated by ELISA. Immunization with pWT, pB.1.351, and INO-4802 induced similar antibody binding titers against the WT and B.1.1.7 variants (**Fig. 2a and c**). Compared to pWT vaccinated animals, there was a small increase in binding titers against P.1 and B.1.351 antigens in mice receiving the pB.1.351 and INO-4802 vaccines. We proceeded to measure the functional ability of the antibodies raised in the vaccinated animals to neutralize SARS-CoV-2. Neutralizing antibody levels against SARS-CoV-2 were measured by a pseudotyped virus neutralization assay in the sera of immunized mice (**Fig. 2b and d**). In the pWT vaccinated animals, while neutralizing activity was similar for B.1.1.7, and P.1 variants as compared to WT, there was a 3-fold reduction in average ID_50_ for the B.1.351 variant. These results are in line with other vaccines matched to the WT spike antigen [7, 28]. In the animals receiving the matched p.B.1.351 vaccine, neutralizing activity was similar for P.1 and B.1.351 variants as compared to WT, and a greater than 7-fold reduction in average ID_50_ for the B.1.1.7 variant. In contrast to the matched vaccines, INO-4802 vaccinated mice demonstrated strong neutralizing activity against all variants assessed. In addition, there was significantly higher neutralizing activity against P.1 and B.1.351 variants in the sera of INO-4802 vaccinated mice, compared to the matched pWT vaccine. Compared to the animals receiving the matched pB.1.351 vaccine, sera from INO-4802 vaccinated animals displayed significantly higher neutralization titers against all the variants tested. Comparison of the sera from INO-4802 vaccinated mice sera to human convalescent sera (HCS) collected in 2020 from donors in the United States demonstrates similar neutralizing activity against the WT and B.1.1.7 variants. While neutralizing activity against the P.1 and B.1.351 variants demonstrates enhanced activity in the INO-4802 vaccinated mice sera compared with HCS (P.1 962 ± 703-1317 vs 279 ± 157-495 and B.1.351 765 ± 552 1058 vs 24 ± 12-46 ID_50_ mean ± range for INO-4802 and HCS, respectively).

**Figure 2.**
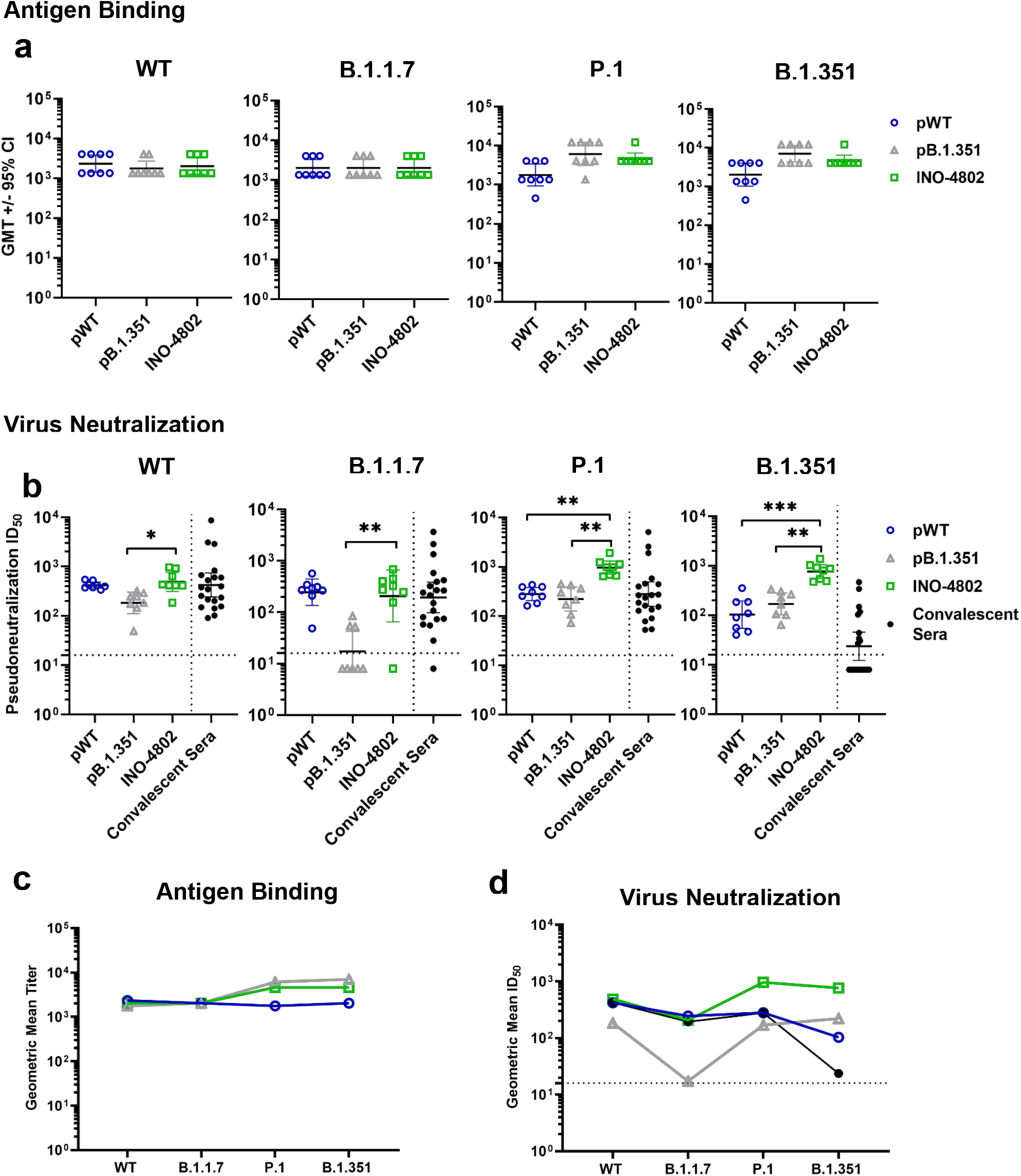
INO-4802 pan-SARS-CoV-2 vaccine-induced humoral immune responses against SARS-CoV-2 VOC. BALB/c mice were immunized on days 0 and 14 with 10µg of pWT, pB.1.351, or INO-4802 and sera samples were collected at day 21 for evaluation of antibody responses as described in the methods. **a)** Sera IgG binding titers against the indicated Spike proteins for pWT, pB.1.351, or INO-4802 vaccinated mice (n of 8 each). Data shown represent geometric mean titer values (GMT+/- 95% CI) for each group of 8 mice. **b)** Sera pseudovirus neutralization ID_50_ titers against the indicated SARS-CoV-2 variant for pWT, pB.1.351, or INO-4802 vaccinated mice (n of 8 per group) or human convalescent sera samples (n of 20). Each data point represents the mean of technical duplicates for individual samples. Dashed lines represent the limit of detection (LOD) of the assay. Samples below LOD were plotted at the number equivalent to half of the lowest serum dilution. *P < 0.05, **P < 0.005, ***P < 0.001 determined by Kruskal-Wallis test (ANOVA) with Dunn multiple comparisons test. **c)** IgG binding data represented as group means for each variant tested. **d)** Pseudovirus neutralization ID_50_ titer data represented as group means for each variant tested.

In summary, INO-4802 demonstrates significantly enhanced neutralizing activity against P.1 and B.1.351 variants while maintaining a strong response against the WT and B.1.1.7 variants, indicating an advantage over variant-matched vaccines (**Fig. 2d**).

### INO-4802 stimulates T cell activity against SARS-CoV-2 Variants

SARS-CoV-2 challenge data in T cell-depleted animals, along with studies indicating reduced disease incidence in individuals harboring pre-existing cross-reactive T cells with SARS-CoV-2 support the importance of COVID-19 vaccine induced cellular immunity [29-31]. We therefore sought to examine T cell responsiveness after INO-4802 vaccination. Splenocytes from mice vaccinated with pWT, pB.1.351, or INO-4802 were stimulated with peptides spanning the WT, B.1.1.7, P.1, and B.1.351 variant Spike proteins. pWT, pB.1.351 and INO-4802 demonstrated induction of comparable T cell responses as measured by IFNγ ELISpot against all variants (**Fig. 3a**). A similar broad T cell reactivity across SARS-CoV-2 VOC was recently reported in human INO-4800 vaccinees [32]. The phenotype of CD4 and CD8 T cell responses was characterized by intracellular cytokine staining on splenocytes isolated from INO-4802-vaccinated mice. Antigen-specific T cells producing IFNγ were observed in both CD4 and CD8 compartments (**Fig. 3b & 3d**). Additionally, CD8 T cells showed expression of CD107a, a marker of cytolytic potential (**Fig. 3c**). The balance of T_H_1 and T_H_2 expressing cells was evaluated based on cytokine expression profile for T_H_1 driving IFNγ and T_H_2 driving IL4 production. CD4 T cells showed greater expression of the canonical T_H_1 cytokine IFNγ relative to IL-4 (**Fig. 3d-f**), consistent with T_H_1-skewed T cell responses following INO-4802 vaccination. Further Th1 vs Th2 evaluation was performed by measuring the induction of IgG2A and IgG1 isotype antibodies. ELISA assay results revealed a higher percentage of IgG2A antibodies compared to IgG1 antibodies in animals vaccinated with pWT and INO-4802, indicative of a T_H_1-biased response (**Supplemental Fig. 2**). Circulating T follicular helper (Tfh) cells are largely representative of a memory CD4 T cell population in the blood that correlates with neutralizing antibody responses, and Tfh cells have been found to be increased in the blood of mice receiving SARS-CoV-2 mRNA vaccines [33-35]. Accordingly, we sought to determine whether Tfh cells were induced following administration of the vaccines. Mice receiving two doses of INO-4802 showed a significantly higher frequency of Tfh cells in circulation relative to mice receiving control pVax (**Fig. 3g**), suggesting enrichment of circulating Tfh cells at an early timepoint post vaccination.

**Figure 3.**
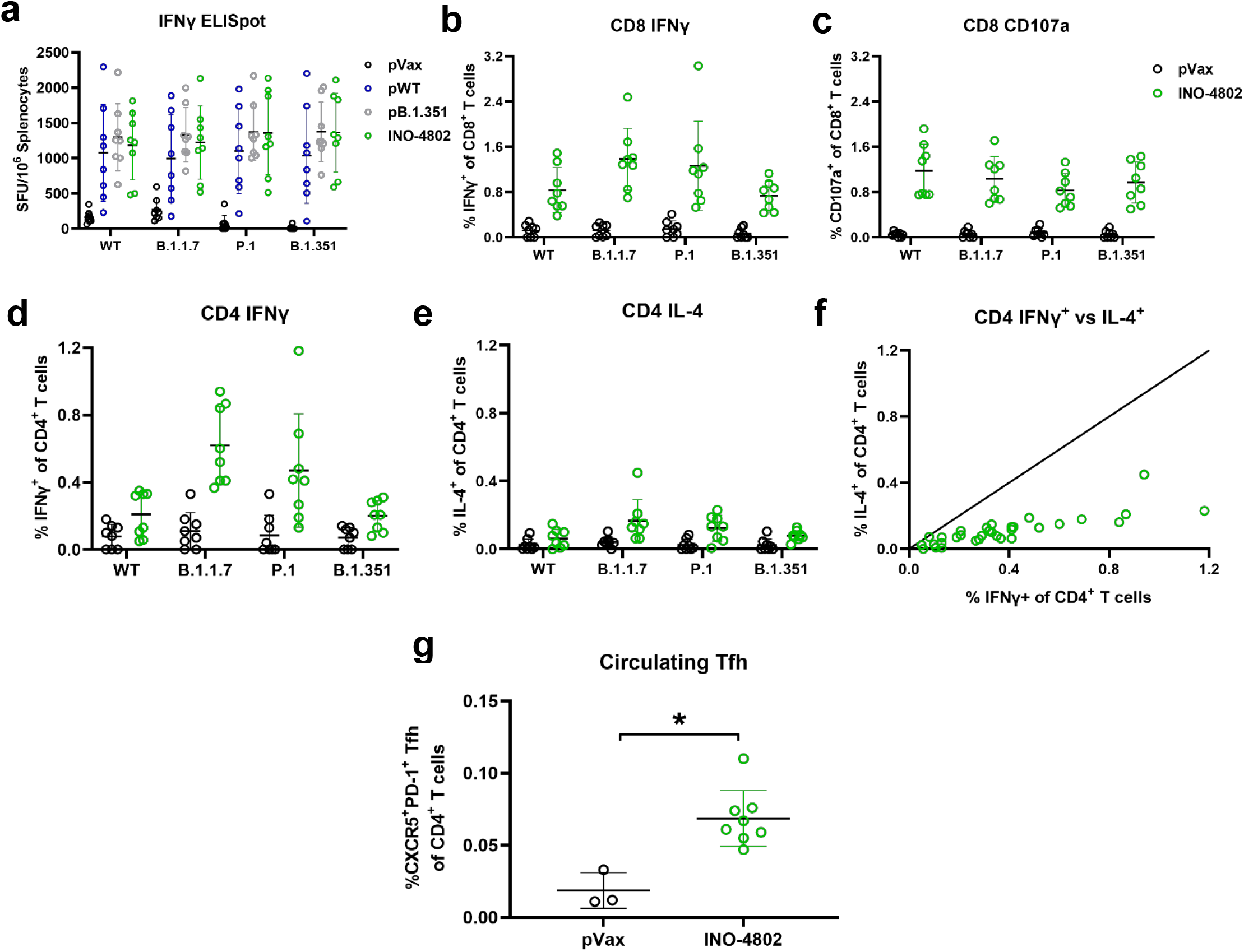
INO-4802 pan-SARS-CoV-2 vaccine-induced cellular immune response against SARS-CoV-2 variants. Splenocytes isolated from mice were collected 1 week after receiving the second dose of either pWT, pB.1.351, or INO-4802. **a)** Splenocytes were stimulated with peptide pools spanning the entire Spike proteins of the WT, B.1.1.7, P.1, or B.1.351 variants and cellular responses were measured by IFNγ ELISpot assay. Mean +/- SD IFNγ SFUs/million splenocytes of experimental triplicates are shown. **b-e)** Intracellular cytokine staining was employed for CD4+ and CD8+ T cell activation. Expression levels of IFNγ, CD107a and IL-4 were analyzed. f) A representative graph showing correlation of T_H_1 (IFNγ) versus T_H_2 (IL-4) cytokine expression in the CD4 compartment of INO-4802 treated animals restimulated with either the WT, B.1.1.7, P.1, or B.1.351 peptide pools. g) Frequencies of circulating Tfh cells (CXCR5^+^PD-1^+^) in CD4 T cells 2 weeks after the second dose of either pWT or INO-4802. *P < 0.05, (Mann-Whitney test).

### INO-4802 confers protection against B.1.351 VOC in Syrian Golden hamsters

The Syrian Golden hamster is permissible to SARS-CoV-2 infection and is the gold standard small animal model for assessing COVID-19 prophylactics [36-40]. We employed this model to test whether the INO-4802 vaccine could confer protection against B.1.351 challenge (**Fig. 4a**). Hamsters receiving two doses of either pB.1.351 or INO-4802 developed high titers of functional antibodies against the B.1.351 variant, as evidenced by strong ACE-2 blocking (**Fig. 4b**) and pseudovirus neutralizing activity (**Fig. 4c**). Spearman correlation was performed between the ACE-2 blocking and pseudovirus neutralization assays and showed significant correlation between assays (r = 0.916, p = 2.77E-06, **Supplemental Fig. 3**) Following challenge, vaccinated hamsters showed only a transient decline in body weight and began to recover from weight loss beyond day 2 post-challenge, while naïve animals continued to decline in body weight until necropsy on day 4 (**Fig. 4d**). Viral titers were undetectable in the lungs of INO-4802-vaccinated hamsters at necropsy (**Fig. 4e**). Lung viral loads were also significantly reduced in hamsters vaccinated with the pWT and pB.1.351-matched constructs (**Fig. 4e**). In addition to the B.1.351 variant, we have assessed the efficacy of INO-4802 protection against WT, B.1.1.7, and P.1 VOCs (**Supplemental Fig. 4**). Currently, against all SARS-CoV-2 VOCs tested we observe maintenance of body weight of INO-4802 vaccinated animals compared to controls.

**Figure 4.**
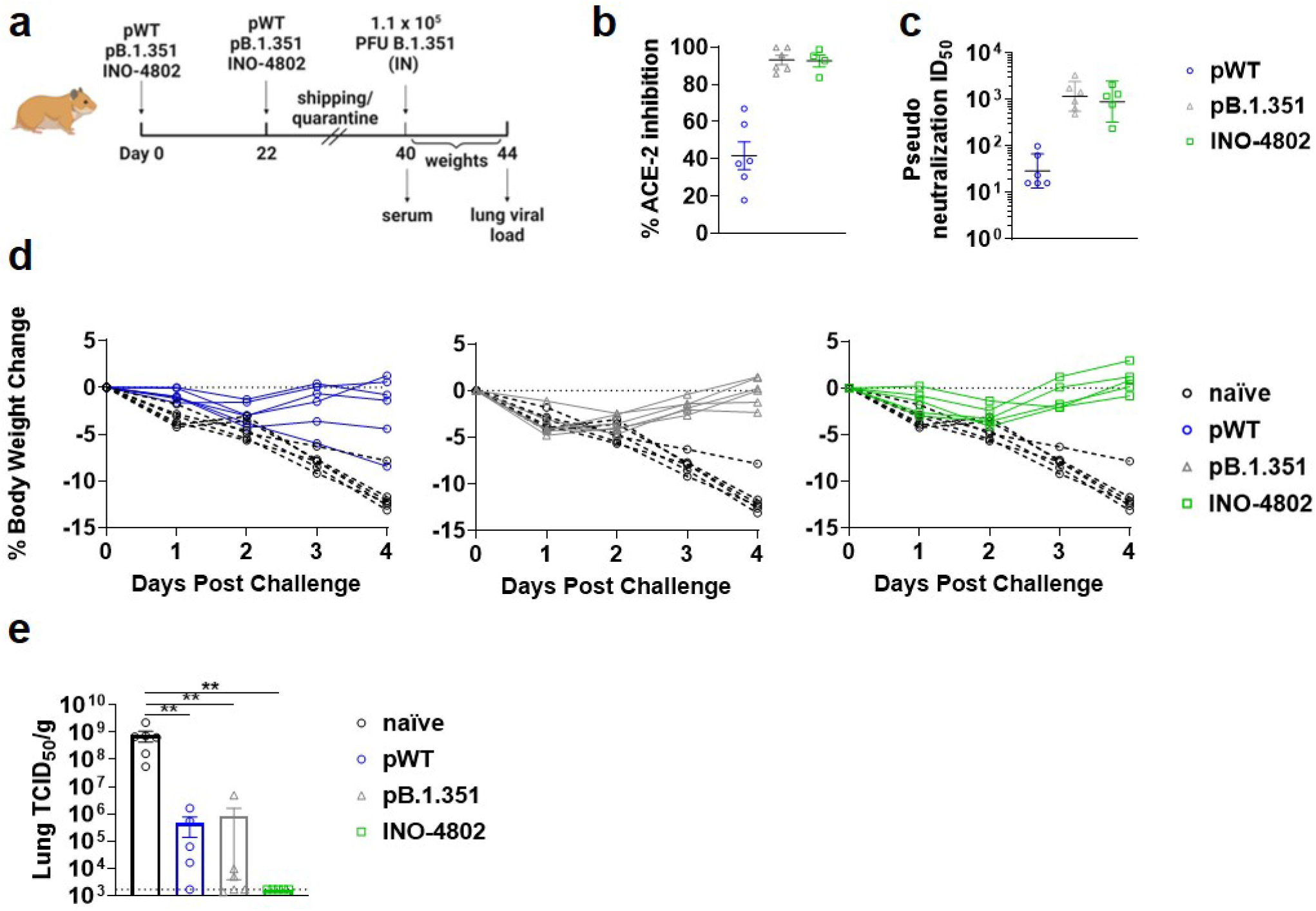
INO-4802 protects Syrian Hamsters against challenge with B.1.351 SARS-CoV-2. **a)** Study schematic – animals received ID+EP immunizations with 95 µg pWT, pB.1.351 or INO-4802 on days 0 and 22. Hamsters were challenged IN with 5.00×10^6 TCID_50_/ 10,875,000 PFU (Bioqual SARS-CoV-2 RSA P4 Lot: 020521-105) and observed for weight loss. On day 4 post challenge animals were euthanized and lung tissue was harvested for viral load measurement. **b)** Sera collected on day 40 were tested for capacity to block binding of human ACE-2 to B.1.351-spike in an electrochemoluminescent-based ELISA assay (mean %inhibition +/-SEM). **c)** Sera collected on day 40 were tested for neutralizing activity in-vitro with B.1.351 pseudovirus (GeoMean ID50 +/- 95%CI). **d)** Weight change of pWT (left, n=6), pB.1.351 (center, n=6) or INO-4802 (right, n=5) vaccinated hamsters compared to unvaccinated animals (n=6) following challenge with B.1.351. **e)** Viral load in the lung at day 4 post challenge as shown by tissue culture ID_50_ (TCID_50_) per gram of lung tissue (mean +/- SEM). Animals with viral load below the LOD of the assay are graphed at 1687 TCID_50_/g, the lower limit of detection. *P < 0.05, **P < 0.005, ***P < 0.001 determined by Mann-Whitney test.

### Heterologous boost with INO-4802 in Syrian hamsters against Global SARS-CoV-2 variants

We employed the hamster model to test whether the INO-4802 vaccine could boost immunity generated by first-generation immunogens based on the WT Spike sequence. Accordingly, we tested the immunogenicity of the pWT in hamsters 236 days after receiving 2 doses of the pWT construct (**Fig. 5a**). Prior to boosting hamsters were split randomly into two groups of four. We compared the scenario of heterologous (INO-4802 construct) to homologous (pWT construct) boost in terms of magnitude and breadth of humoral responses targeting the VOCs. VOC antigen binding titers were increased across all variants tested (**Fig. 5b**). We observed enhanced levels in the INO-4802 group compared to the homologous boost group (pWT GMTs 21,044 and INO-4802 109,350). As the pre-boost levels in the INO-4802 group were slightly higher than the homologous group we assessed the fold changes in binding titers between groups. We observed a trend towards higher log2-fold increase of endpoint binding titers after boost with INO-4802 (3.6 4.4 log2-fold change) compared to boost with pWT (2.0 – 2.4 log2 fold change). These results suggested broad humoral responses against global SARS-CoV-2 variants that were boosted with the pan-SARS-CoV-2 construct compared to the homologous construct.

**Figure 5.**
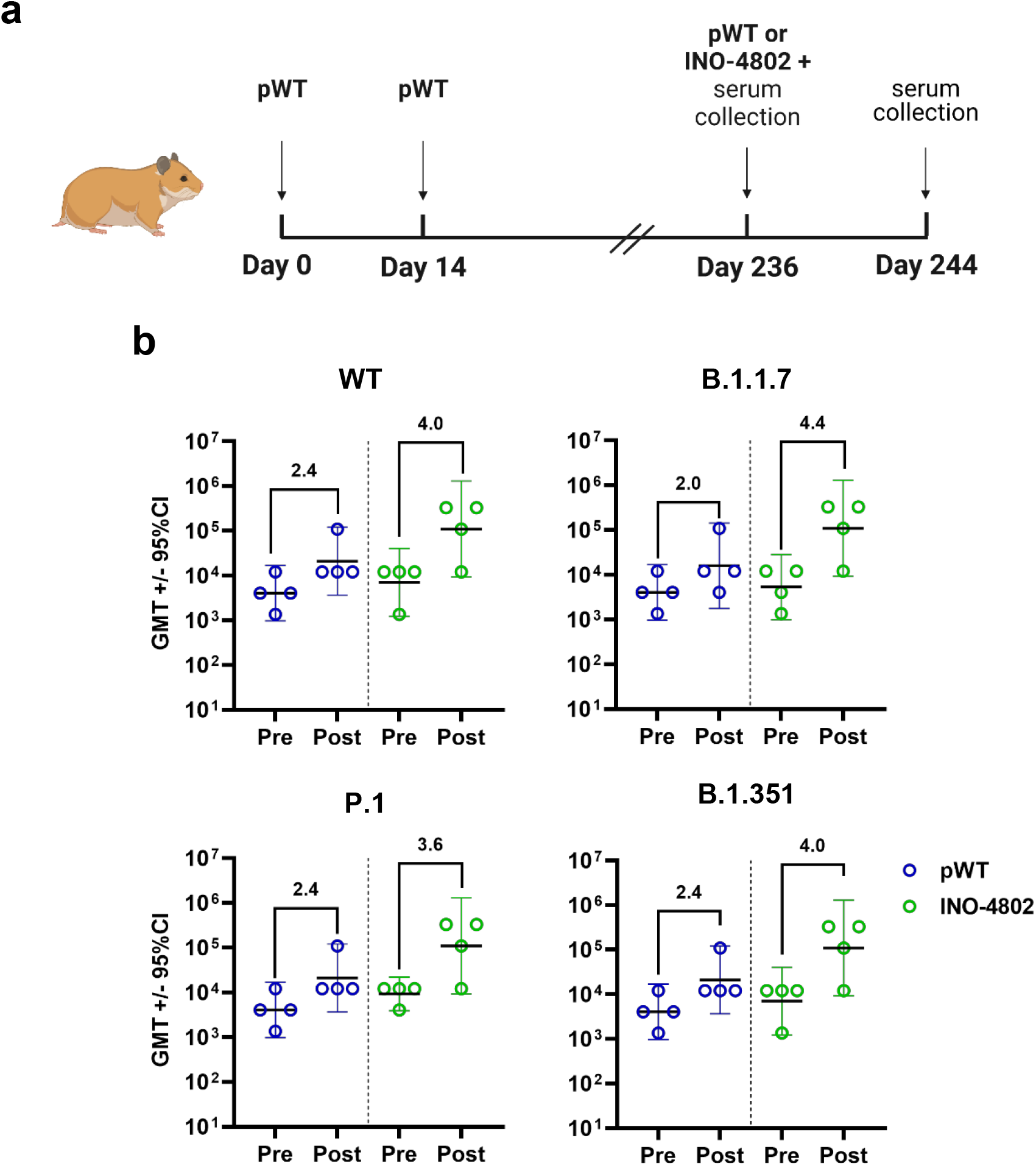
Heterologous boost with INO-4802 pan-SARS-CoV-2 vaccine induces humoral immune responses against SARS-CoV-2 variants in Syrian Hamsters. **a)** Experimental design. Syrian hamsters (n=8) were immunized with 90 µg of pWT on days 0 and 14. On day 236 animals were boosted with 90 µg of pWT or INO-4802 **b)** Pre- and post-boost sera IgG binding titers against the indicated SARS-CoV-2 Spike antigens. Symbols represent endpoint binding titers for individual animals and lines and bars represent group GMT +/- 95%CI. Values indicate log2 fold changes of GMTs from pre- to post- boost.

## Discussion

Here we present the preclinical immunogenicity results for a pan-SARS-CoV-2 DNA vaccine, INO-4802 designed with SynCon technology to provide broad immune responses against SARS-CoV-2 Spike antigen on emerging VOC. INO-4802 induced broadly neutralizing antibodies and T cell responses against WT, B.1.1.7, P.1, and B.1.351 SARS-CoV-2 Spike variants in BALB/c mice. In contrast, the cross-neutralizing activity for strain matched vaccines (pWT and pB.1.351) were limited. Importantly, hamsters vaccinated with INO-4802 were completely protected from challenge with B.1.351. Initial data demonstrates the potential of INO-4802 as a pan-SARS-CoV-2 vaccine countermeasure to emerging VOC.

Current COVID-19 vaccines authorized for emergency use in the US were designed early in the pandemic to match the initial outbreak SARS-CoV-2 strain. The natural mutation rate in RNA viruses inevitably spurred selection for new variants and accumulating evidence points to some critical variants showing enhanced spread, ability to escape known neutralizing antibodies, or leading to higher hospitalization rates [10, 41, 42]. Vaccine trials have already reported reduced efficacy in geographical regions in which the B.1.351 VOC is present [13, 43, 44]. In response, vaccines matched to B.1.351 have been designed [45]. In addition to the pan-SARS-CoV-2 vaccine INO-4802, we also tested a DNA vaccine matched to the B.1.351 VOC as a comparator matched VOC strain design. However, while we observed immune responses to the matched antigen, functional antibody responses to other tested variants were reduced (**Fig. 2b**). These data indicate that focusing on single variant, matched strain approach is to some extent effective but may limit cross-protective potential against diverse novel VOCs, at worst becoming obsolete if facing rapid variant shifts during the development process. This concern is not theoretical. Nearly simultaneous occurrences of B.1.1.7 and B.1.351, followed by P.1 a short time later, show how rapidly the variant landscape can shift. Recent SARS-CoV-2 infection rate surges in India demonstrate the speed at which changes can occur [46]. In response, we have initiated testing of INO-4802-induced immunity against the B.1.617.2 (Delta) variant (**Supplemental Fig. 5**). Initial data indicates functional antibodies blocking ACE-2 binding to the B.1.617.2 spike are induced and protection against viral challenge is achieved by INO-4802 vaccination. This finding is encouraging and supportive of the SynCon approach as B.1.617.2 has recently been identified as a VOC and was not included in the design of INO-4802 immunogen. Future studies will test whether activity is maintained against other VOCs which emerge.

In the design process of INO-4802, we utilized a broad antigen design strategy using SynCon technology as a potential mitigation solution to the limited coverage provided by a matched strain approach. A single construct design consisting of an antigen representative of multiple viral strains while retaining sufficient identity to generate effective and broad immunity is a favorable solution that presents fewer production barriers as defined by cost of goods and formulation complexity. Application of the SynCon antigen design platform in conjunction with rational design choices allowed us to generate a pan-SARS-CoV-2 vaccine design, INO-4802 (**Fig. 1a, b**). We evaluated this approach in terms of the broadness of the immune response induced by INO-4802 against the VOC.

As detailed in this study, humoral immunogenicity results in BALB/c mice demonstrated the consensus-based approach provided broad cross-reactive neutralizing antibody responses against all the VOC tested (**Fig. 2b**). While the matched pWT construct and INO-4802 performed similarly in neutralization assays against WT and B.1.1.7-matched pseudotyped virus, strain-matched vaccination with pB.1.351 construct showed poor overall performance against B.1.1.7. This may be due to the unique cluster of mutations in the B.1.351 Spike compared to either WT or to INO-4802, making it a less ideal antigen for promoting a B.1.1.7-directed humoral response.

B.1.351 has multiple unique changes to the N-terminal domain (NTD) which along with the RBD contains potent neutralization sites [47]. Generation of neutralizing response to B.1.351 alone may come at the cost of a lack of neutralizing antibody response to more diverse lineages. INO-4802 demonstrated superiority to both pWT and pB.1.351 in inducing functional antibodies against P.1 and B.1.351-matched pseudoviruses (**Fig. 2d**). The greater magnitude of neutralizing activity induced by INO-4802 in BALB/c mice compared to pB.1.351 against P.1 VOC may be less surprising than those against B.1.351 VOC, as the pB.1.351 was strain-matched in this case. Additionally, a trend for improved protection against B.1.351 challenge was observed in INO-4802 vaccinated hamsters compared to animals receiving the strain matched pB.1.351 vaccine (**Fig. 4**). Added design features in INO-4802 to promote antigen stability, notably the 2P mutation, may be an important structural advantage to account for this observed difference.

Vaccination with pWT, pB.1.351 and INO-4802 resulted in comparably strong IFNγ ELISpot responses against WT and VOC-matched peptide pools in a murine model (**Fig. 3**). Lack of differentiation between the vaccine constructs in respect to level of T cell immunity against VOC Spike antigens was expected. The highly diverse and linear epitope dependent T cell compartment is less impacted than the structurally dependent functional antibody response. The results are supported by the maintenance of pWT T cell immunity against the same panel of VOC [32], as well as data in COVID-19 exposed donors and vaccinees showing a negligible impact of SARS-CoV-2 variants on CD4 and CD8 T cell responses [48]. In addition to ELISpot analysis, we performed additional functional T cell assays. INO-4802 vaccination induced CD107a/IFN-γ+ cross-reactive CD8 T cells and Tfh cells, cell populations involved in cytotoxic and augmenting B cell responses, respectively (**Fig. 3b & c**). Importantly, Inovio’s plasmid-encoded antigens and *in vivo* electroporation technology have repeatedly been shown to drive cellular immune responses in humans [49-51]. The cross-reactive T cell results may also play an important role in effective cross-protection, in a similar manner to that observed in individuals during the H1N1 influenza pandemic. In a study of healthy individuals who contracted pH1N1 influenza during the 2009 pandemic, the presence of pre-existing T cells responsive to conserved epitopes in pH1N1 correlated with less severe clinical outcomes even when effective cross-neutralizing antibodies were not present [52]. Support for an important role of T cells was provided in the SARS-CoV-2 challenge study (**Fig. 4**). In this B.1.351 SARS-CoV-2 challenge study we observed significant protection against both weight loss and reduction in lung viral loads in animals immunized with the pWT vaccine (**Fig. 4**). In contrast to the decline in neutralizing antibodies in pWT immunized animals against B.1.351, the level of T cell activity against the VOCs was fully maintained (**Fig. 3**). Such observations support the role of the T cell compartment in providing protection against COVID-19 disease.

In addition to the use of INO-4802 as a prime vaccine, we assessed employing it in a heterologous boost regimen (**Fig. 5a**). INO-4802 boosting effect was measured approximately 8 months after priming with pWT vaccine in the Syrian Golden hamster model. Delaying the INO-4802 boost by approximately 8 months provide time for the maturation of the immune response the first round of vaccination and potentially antigenic imprinting to WT spike antigen. However, initial humoral immunogenicity readout suggests strong boost of binding antibody titers across the panel of WT and VOC antigens. The increase in binding titers against all the VOCs tested was greater than same dose boost of pWT. Further studies are ongoing and planned to fully investigate INO-4802 as a boost to first generation vaccine responses generated by INO-4800 and other platform technologies.

We have reported on the design and testing of a pan-SARS-CoV-2 vaccine, INO-4802. High levels of cross-reactive neutralizing activity against multiple VOCs was demonstrated by pseudotyped virus neutralization, and superior protection against B.1.351 challenge was observed compared to a strain-matched vaccine (**Fig. 4e**). Additionally, INO-4802 has shown utility in both prime and heterologous boost immunization scenarios. Vaccination with INO-4802 led to cross-reactive T cell immune responses as well as circulating populations of Tfh cells correlative of both memory and neutralizing antibody responses. In summary, INO-4802 represents a potentially useful pan-SARS-CoV-2 vaccine approach against multiple variants of concern and the data presented in this report supports its further development.

## Methods

### Design and generation of plasmid constructs

The synthetic consensus (SynCon®) sequence for the SARS-CoV-2 spike glycoprotein with additional focused RBD changes, dual proline mutations (K986P/V987P (2P)) and the endogenous SARS-CoV-2 Spike signal peptide replaced with IgE leader sequence was codon-optimized using Inovio’s proprietary optimization algorithm. A Kozak sequence (GCCACC) immediately 5’ of the start codon in addition to restriction sites for subcloning of the construct into pGX0001 vector (5’ BamHI and 3’ XhoI) were added. The optimized DNA sequence was synthesized (Genscript, Piscataway NJ), digested with BamHI and XhoI, and cloned into the expression vector under the control of the cytomegalovirus immediate-early promoter, generating INO-4802. Strain-matched spike sequences (pWT, pB.1.351) were similarly optimized and cloned into identical restriction site locations into the pGX0001 backbone. Pseudovirus plasmids were designed and constructed as previously described [32]. Sequence analysis was performed using custom Python scripts (Python Software Foundation, https://www.python.org/) and assembly was performed using Geneious Prime® 2020.2.3 (Build 2020-08-25, Biomatters Ltd., Auckland NZ). Molecular modeling of the spike was performed with multiple SARS-CoV-2 spike templates (PDB ID:6XM3, 6VXX, 7K4N) using Prime [Jacobson 2004] from the Bioluminate suite (BioLuminate Release 2018-3, Schrödinger, LLC, New York, NY). Visualization was performed using Bioluminate and Discovery Studio (Discovery Studio 2019, BIOVIA, Dassault Systèmes, San Diego, CA).

### Cell lines and *In-vitro* plasmid expression

HEK-293T (ATCC® CRL-3216™) and African Green monkey kidney COS-7 (ATCC® CRL-1651™) cell lines were obtained from ATCC (Old Town Manassas, VA). All cell lines were maintained in DMEM supplemented with 10% fetal bovine serum (FBS) and penicillin-streptomycin.

For in vitro protein expression by Western blot, human embryonic kidney cells, 5.5×10^5^ 293T were transfected with 2.5µg pDNA in 6-well plates using Lipofectamine 3000 (Invitrogen #L3000015) transfection reagent following the manufacturer’s protocol. Transfections were performed in duplicate wells for each plasmid DNA. Forty-eight hours later cell lysates were harvested using Cell Signaling Cell Lysis Buffer (#9803) and duplicate transfection lysates pooled for expression analysis. Proteins were separated on a 4–12% BIS-TRIS gel (ThermoFisher Scientific), then following transfer, blots were incubated with an anti-SARS-CoV spike protein polyclonal antibodies (S1, Sino Biological #40591-T62; S2, Invitrogen #PA1-41165; RBD, Sino Biological #40592-MP01) then visualized with horseradish peroxidase (HRP)-conjugated anti-mouse or anti-rabbit IgG (Bethyl) (GE Amersham). Beta actin was detected using Santa Cruz # SC-47778.

For *in vitro* RNA expression by qRT-PCR, transfections, RNA purification, cDNA synthesis, and qPCR assay were performed as previously described [20, 32]. PCR was performed using a single set of primers and probes recognizing the RNA products of all three plasmids (pS-spike forward ATGATCGCCCAGTACACATC, pS-spike reverse CACGCCGATGCCATTAAATC, pS-spike probe AT CACCAGTGGCTGGACATTTGGA). In a separate reaction, the same quantity of sample cDNA was subjected to PCR using primers and a probe designed (β-actin Forward – GTGACGTGGACATCCGTA AA; β-actin Reverse – CAGGGCAGTAATCTCCTTCTG; β-actin Probe – TACCCTGGCATTGCTGACAGGATG) for COS-7 cell line β-actin sequences. The primers and probes were synthesized by Integrated DNA Technologies, Inc. and the probes were labeled with 56-FAM and Black Hole Quencher 1.

### *In-vivo* immunogenicity

BALB/c mice (6 weeks old, Jackson Laboratory, Bar Harbor, ME) and Syrian Golden Hamsters (8 weeks old, Envigo, Indianapolis, IN) were housed at Acculab (San Diego, CA). Mice (n=8/group) received 30 µL intramuscular (IM) injection of 10 µg pDNA immediately followed by electroporation (EP) in the tibialis anterior (TA) muscle on days 0 and 14 of the experiment. Hamsters (n=4/group) received 60 µL IM injection of 90 µg pDNA immediately followed by EP into the TA on days 0, 14 and 236 of the experiment. The CELLECTRA® EP treatment consists of two sets of pulses with 0.2 Amp constant current. Second pulse set is delayed 4 s. Within each set there are two 52 ms pulses with a 198 ms delay between the pulses. Mice were euthanized on day 21 for terminal blood collection and spleens were harvested for cellular assays. Serum was collected from hamsters on days 236 (pre-boost) and 244 (post-boost) by jugular blood collection for pseudovirus-neutralization assay. All animal treatments and procedures were performed at Acculab, and animal testing and research complied with all relevant ethical regulations and studies received ethical approval by the Acculab Institutional Animal Care and Use Committees (IACUC).

### B.1.351 SARS-CoV-2 challenge study

Syrian golden hamsters (8 weeks old, Envigo, Indianapolis, IN) were housed at Acculab (San Diego, CA). For the SARS-CoV-2 challenge study, hamsters (n=6/group) received 100 µL ID injection of 95 µg pDNA immediately followed by EP as described above into the flank on days 0 and 22 of the experiment. Following treatments at Acculab, animals were transported to Bioqual Inc. (Rockville, MD) for challenge. 40 days following the initial pDNA treatment, intranasal inoculation (IN) was performed on Ketamine/Xylazine anesthetized hamsters. The animals were challenged with 5.00×10^6 TCID_50_/ 10,875,000 PFU (Bioqual SARS-CoV-2 RSA P4 Lot: 020521-105), with a total volume of 100 μL per animal (50μL/nostril). Post-challenge, the animals were weighed daily, beginning the day of challenge. Serum was collected from hamsters on days 40 (pre-challenge) by jugular blood collection for pseudovirus-neutralization assay. All animal testing and research complied with all relevant ethical regulations and studies received ethical approval by the Acculab and Bioqual Institutional Animal Care and Use Committees (IACUC).

### TCID_50_ Assay for Determination of Infectious Viral Load

For infectious titer determination from lungs samples, TCID_50_ assay was performed on the lung tissues harvested from the hamsters. To set up the assay, frozen lung tissue was placed in 15 mL conical tube on wet ice containing 0.5 mL media and homogenized for 10-30 secs (Probe, Omni International: 32750H). The tissue homogenate was spun to remove debris at 2000g, 4°C for 10 min. The supernatant was passed through a strainer that was placed on original vial, placing vials on wet ice. 20 μL of this supernatant was tested in the assay in quadruplicate in a 96 well plate format.

To perform this assay, Vero TMPRSS2 cells are plated at 25,000 cells per well in DMEM + 10% FBS + Gentamicin. The plate was incubated at 37°C, 5.0% CO2. The cells should be 80 -100% confluent the following day. When the 80-100% is confirmed, the media was aspirated out and replaced with 180μL of DMEM + 2% FBS + gentamicin. Then 20 μL of the sample was added to top row in quadruplicate. The top row was mixed 5 times with a pipette and titer down 20 μL, representing 10-fold dilutions. The pipette tips were disposed of between each row and the mixing is repeated until the last row on the plate. The plates for the samples are incubated again at 37°C, 5.0% CO_2_ for 4 days. After 4 days, the plates were visually inspected for CPE. Non-infected wells would be expected to have a clear confluent cell layer, unlike infected cells, which would demonstrate cell rounding. The presence of CPE was recorded as a plus (+) and absence of CPE as minus (-). The TCID_50_ was then calculated using the Read-Muench formula.

### Antigen binding Assays

Binding ELISAs were performed as described previously [32] except different variants of SARS-CoV-2 S1+S2 spike proteins were used for plate coating. Binding titers were determined after background subtraction of animals vaccinated with mock vector. The S1+S2 wild-type spike protein (Acro Biosystems #SPN-C52H8) contained amino acids 16-1213 of the full spike protein (Accession #QHD43416.1) with F817P, A892P, A899P, A942P, K986P, V987P, R683A and R685A mutations to stabilize the trimeric prefusion state. The B.1.1.7 and B.1.351, and P.1 S1+S2 variant proteins (Acro Biosystems #SPN-C52H6, #SPN-C52Hc, and #SPN-C52Hg, respectively) additionally contained the following proline substitutions for trimeric protein stabilization: F817P, A892P, A899P, A942P, K986P, and V987P. The B.1.1.7 protein contained the following variant-specific amino acid substitutions: HV69-70del, Y144del, N501Y, A570D, D614G, P681H, T716I, S982A, D1118H; and the B.1.351 protein contained the following substitutions: L18F, D80A, D215G, R246I, K417N, E484K, N501Y, D614G, A701V; and the P.1 protein contained the following:L18F,T20N,P26S,138Y,R190S,K417T,E484K,N501Y,D614G,H655Y,T1027I,V1176F. Half-area assay plates were coated using 25 µL of 1 µg/mL of protein. Secondary antibodies included IgG (Sigma #A4416), IgG2A (Abcam #ab98698), and IgG1 (Abcam #ab98693) at 1:10,000 dilution.

### Human ACE-2 blocking assay

Serum was tested for blocking of ACE-2 binding with V-Plex COVID-19 Ace2 Neutralization Panel V Kit (MSD). The manufacturers procedure was followed. Briefly, 150 µl/well of Blocker A solution was added to the assay plate. Plate was sealed and incubated for one hour at room temperature with shaking (700 rpm). Sera were diluted 1:30 dilution. Diluent alone was used as a blank in the calculations. Assay plate was then washed 3 times with 200 µL/well of 1x MSD wash buffer. 25 µL/well of diluted samples were added to the assay plate in duplicates. The assay plate was sealed and incubated for one hour at room temperature with shaking (700 rpm). Sulfo-Tag ACE2 Protein reagent was diluted 1:200 and 25 µL/well was added to each well of the plate. The assay plate was sealed and incubated for one hour at room temperature with shaking (700 rpm) and then washed 3 times with 200 µL/well of 1x MSD wash buffer. 150 µL/well of MSD Gold Read Buffer was added to each well, and plate was immediately read on Meso Sector 600 machine. Data were analyzed using MSD Discovery Workbench Software and %inhibition was calculated as %inhibition = (1 – (average sample ECL signal/average ECL signal of diluent only)) x 100.

### Pseudovirus production

SARS-CoV-2 pseudotyped stocks encoding for the WT, B.1.1.7, P.1, or B.1.351 Spike protein (**Table 2**) were produced using HEK293T cells transfected with Lipofectamine 3000 (ThermoFisher) using IgE-SARS-CoV-2 S plasmid variants (Genscript) co-transfected with pNL4-3.Luc.R-E-plasmid (NIH AIDS reagent) at a 1:8 ratio. Cell supernatants containing pseudotyped viruses were harvested after 72h, steri-filtered (Millipore Sigma), and aliquoted for storage at - 80°C.

**Table 2.** Spike glycoprotein mutations in antigens.

| Pseudoviral construct | Mutations |
| --- | --- |
| WT | Original Wuhan sequence (NC_045512) |
| B.1.1.7 | Δ69-70/Δ144/N501Y/A570D/D614G/P681H/T716I/S982A/D1118H |
| P.1 | L18F/T20N/P26S/D138Y/R190S/K417/E484K/N501Y/D614G/H655Y/T1027I/V1176F |
| B.1.351 | L18F/D80A/D215G/Δ242-244/R246I/K417N/E484K/N501Y/D614G/A701V |

### SARS-CoV-2 Pseudotyped neutralization

CHO cells stably expressing ACE2 (ACE2-CHOs) to allow permissiveness to SARS-CoV-2, were seeded at 10,000 cells/well. SARS-CoV-2 pseudotyped stocks were titered to yield greater than 30 times the cell only control relative luminescence units (RLU) 72h post-infection. Sera from vaccinated mice were heat inactivated and serially diluted two-folds starting at 1:16 dilution. Sera were incubated with SARS-CoV-2 pseudotyped virus for 90 min at room temperature. After incubation, sera-pseudovirus mixture was added to ACE2-CHOs and allowed to incubate in a standard incubator (37°C, 5% CO_2_) for 72h. After 72h, cells were lysed using Bright-Glo™ Luciferase Assay (Promega) and RLU was measured using an automated luminometer. Neutralization titers (ID_50_) were calculated using GraphPad Prism 8 and defined as the reciprocal serum dilution at which RLU were reduced by 50% compared to RLU in virus control wells after subtraction of background RLU in cell control wells.

### COVID-19 Convalescent Serum samples

Two sets of convalescent donor sera were used in this study, each consisting of 10 donors from the USA. One set included donors that tested positive for SARS-CoV-2 infection in March-April 2020, while the other set consisted donors that tested positive in October 2020. Samples were collected between 14 and 71 days from the onset of symptoms. 15 donors (75%) had disease symptoms classified as mild, 1 donor (5%) was asymptomatic, and 4 donors (20%) had moderate disease symptoms. Seven donors were male, and thirteen were female. Ages ranged from 19 to 60 years. Serum samples were sourced from BioIVT.

### SARS-CoV-2 spike ELISpot assay

Mouse Peripheral mononuclear cells (PBMCs) post-vaccination with DNA vaccinations were stimulated *in vitro* with 15-mer peptides (overlapping by 9 amino acids) spanning the full-length Spike protein sequence of the indicated variants. Variant peptide pools (GenScript, custom) included the following changes to match published deletions/mutation in each variant: B.1.1.7 variant (delta69-70, delta144, N501Y, A570D, D614G, P681H, T716I, S982A, D1118H), P.1 variant L18F, T20N, P26S, D138Y, R190S, K417T, E484K, N501Y, D614G, H655Y, V1176F), B.1.351 variant (L18F, D80A, D215G, delta242-244, R246I, K417N, E484K, N501Y, D614G, A701V); Cells were incubated overnight in an incubator with peptide pools at a concentration of 1 μg per ml per peptide in a precoated ELISpot plate, (MabTech, Mouse IFNγ ELISpot Plus). Cells were washed off, and the plates were developed via a biotinylated anti-IFN-γ detection antibody followed by a streptavidin-enzyme conjugate resulting in visible spots. Each spot corresponds to an individual cytokine-secreting cell. After plates were developed, spots were scanned and quantified using the CTL S6 Micro Analyzer (CTL) with *ImmunoCapture* and *ImmunoSpot* software. Values are shown as the background-subtracted average of measured triplicates. The ELISpot assay qualification determined that 12 spot forming units was the lower limit of detection. Thus, anything above this cutoff is considered to be a signal of an antigen specific cellular response.

### INO-4802 SARS-CoV-2 spike flow cytometry assays

Mouse splenocytes were also used for intracellular cytokine staining (ICS) analysis and visualized using flow cytometry. One million splenocytes in 200 µL complete RPMI media were stimulated for six hours (37 °C, 5% CO_2_) with DMSO (negative control), PMA and ionomycin (positive control, 100 ng/mL and 2 μg/mL, respectively), or with the indicated peptide pools (225 ug/mL). Variant peptide pools included the following changes to match published deletions/mutation in each variant: B.1.1.7 variant (delta69-70, delta144, N501Y, A570D, D614G, P681H, T716I, S982A, D1118H), P.1 variant L18F, T20N, P26S, D138Y, R190S, K417T, E484K, N501Y, D614G, H655Y, V1176F), B.1.351 variant (L18F, D80A, D215G, K417N, E484K, N501Y, D614G, A701V).

After stimulation, cells were washed in PBS for live/dead staining (Life Technologies Live/Dead aqua fixable viability dye), and then stained with extracellular markers. The cells were then fixed and permeabilized (eBioscience™ Intracellular Fixation and Permeabilization Buffer Set) and then stained for the indicated intracellular cytokines using fluorescently-conjugated antibodies (**Table S1**). In a separate flow cytometry assay, circulating T follicular helper (Tfh) cells were assessed in vaccinated mice using a whole blood staining strategy. 100 uL of whole blood was obtained 2 weeks after the second dose of either INO-4802 or pVax. Whole blood was directly stained using the same viability dye and fluorescently-conjugated antibodies described above for CD3, CD4, and CD8. The antibody cocktail also included the canonical Tfh markers CXCR5-biotin (BD Biosciences), PD-1 PE-CF594 (BD Biosciences), and ICOS BV650 (BD Biosciences) in the presence of Fc block (BD Biosciences). Following a 45-minute incubation at 4 C, whole blood was lysed using FACS Lysing Solution (BD Biosciences) according to the manufacturer’s instructions. Cells were then stained with streptavidin-BV421 (BD Biosciences) for 40 minutes at 4°C, fixed, and acquired on a FACSCelesta flow cytometer (BD Biosciences). Data were analyzed using FlowJo software. Tfh cells were identified as CD4^+^CXCR5^+^PD-1^+^ T cells.

### Data analysis

GraphPad Prism 8.1.2 (GraphPad Software, San Diego, USA) was used for graphical and statistical analysis of data sets. P values of < 0.05 were considered statistically significant. A nonparametric Mann-Whitney test was used to assess statistical significance when comparing two groups and a by Kruskal-Wallis test (ANOVA) with Dunn multiple comparisons test when comparing three or more groups.

## Data availability

The data that support the findings of this study are available from the corresponding authors upon reasonable request.

## Acknowledgments

We acknowledge the members of the Inovio Pharmaceuticals Vivarium Ops team, Olivia Bedoya, Ariza Delgado, Gloria Mendez and Jon Schantz, for significant technical assistance. We acknowledge GISAID and the network of researchers who share valuable sequencing data. This work is funded by Coalition for Epidemic Preparedness Innovations (CEPI).

## Author contributions

Conceptualization K.E.B. J.J.K., D.B.W., L.M.H., V.M.A., D.E., R.K., S.J.R., C.C.R., K.S., T.R.F.S.; Methodology V.M.A., R.K., S.J.R., C.C.R., K.S., T.R.F.S., J.W., M.Y., J.F, R.F. ; Investigation V.M.A., K.S., D.E., J.N.W., B.S., I.M., M.V., A.D., M.V., Z.E., P.P., D.A., S.K, M.P, B.N, M.L, B.S, H.B, H.H; Resources K.E.B. J.J.K., L.M.H.; Writing-original draft preparation V.M.A., D.E., S.J.R., C.C.R., B.S., K.S., T.R.F.S., J.T., J.W.; Writing-review and editing K.E.B., J.J.K., D.B.W., L.M.H., S.J.R., C.C.R., T.R.F.S., J.T.; Project administration D.B.W., S.J.R., J.J.K., L.M.H., T.R.F.S., K.E.B.

## Competing interests

C.C.R., V.M.A., R.K., K.S., J.T., B.S., D.E., J.W., I.M., A.D., Z.E., P.P., D.A. M.Y., J.F., R.F., M.V., J.J.K., L.M.H., S.J.R., T.R.F.S., K.E.B. are employees of Inovio Pharmaceuticals and as such receive salary and benefits, including ownership of stock and stock options, from the company. D.B.W. has received grant funding, participates in industry collaborations, has received speaking honoraria, and has received fees for consulting, including serving on scientific review committees and board services. Remuneration received by D.B.W. includes direct payments or stock or stock options, and in the interest of disclosure he notes potential conflicts associated with this work with Inovio and possibly others. In addition, he has a patent DNA vaccine delivery pending to Inovio. All other authors report there are no competing interests.

## Additional Information

Supplementary information is available for this paper at (Link)

**Table S1.** Flow cytometry panel.

| Tube | Channel | Marker/Cytokine |
| --- | --- | --- |
| 1 | FITC | CD107a |
| 2 | V500 | Live/Dead Fix Aqua |
| 3 | PerCP-Cy5.5 | CD4 |
| 4 | BV786 | CD8 |
| 5 | APC-Cy7 | CD3 |
| 6 | BV605 | IFNγ |
| 7 | AlexaFluor 700 | IL-2 |
| 8 | AlexaFluor647 | IL-4 |
| 9 | V450 | TNF |

**Supplemental Figure 1.**
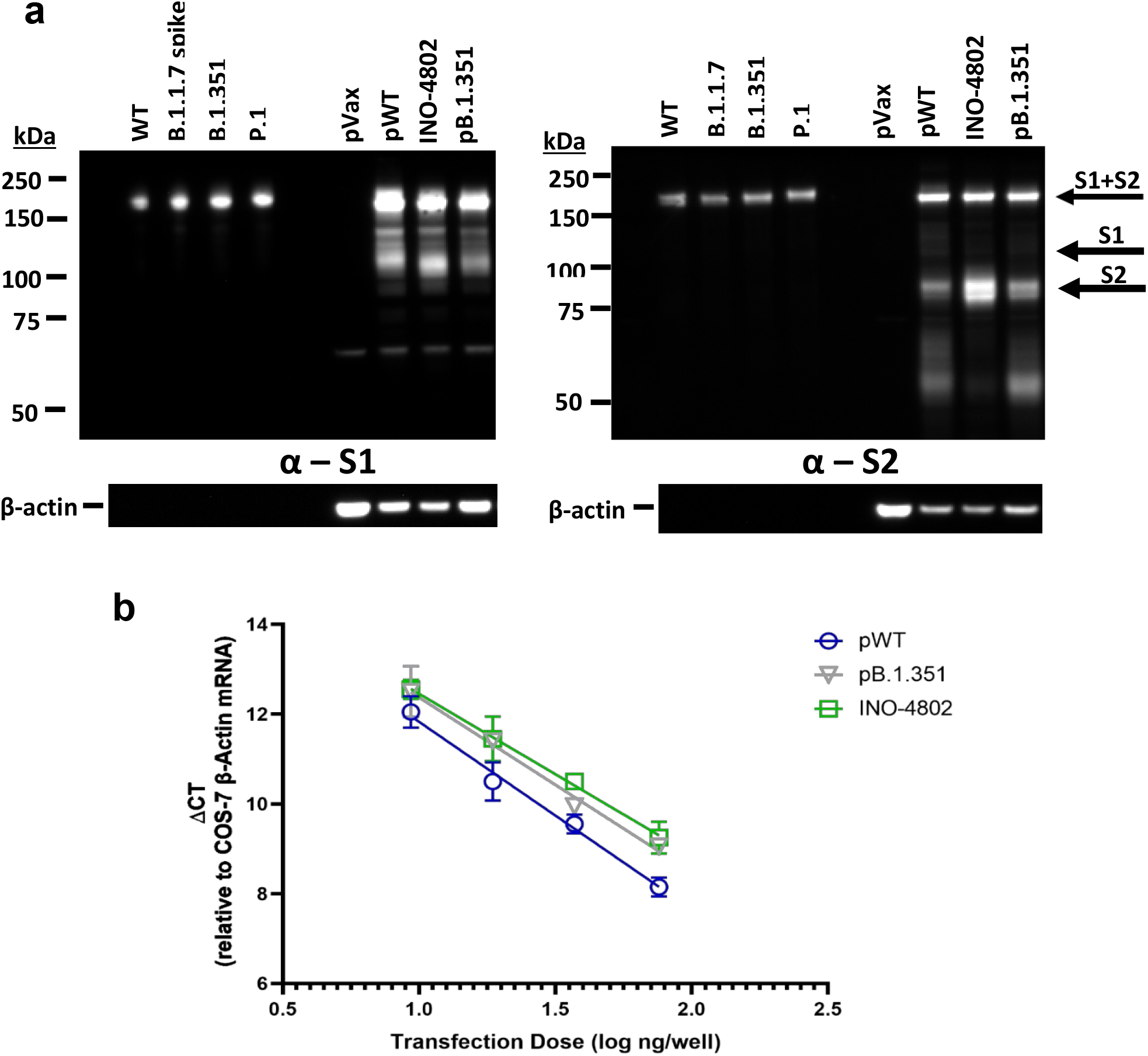
*In vitro* expression of pDNA. **a)** Analysis of *in vitro* expression of Spike protein after transfection of HEK-293T cells with empty vector (pVax), pWT, INO-4802, or pB.1.351 plasmid by Western blot. Control proteins and HEK-293T cell lysates were resolved on a gel and probed with a polyclonal anti-SARS-CoV-2 S1 (left) and S2 protein (right). Blots were stripped then probed with an anti-β-actin loading control. **b)** *In vitro* expression of RNA by RT-PCR assay. RNA extracts from COS-7 cells transfected in duplicate with pWT, INO-4802, or pB.1.351. Extracted RNA was analyzed by RT-PCR using a PCR assays designed for SARS-CoV-2 spike and for COS-7 β-Actin mRNA, used as an internal expression normalization gene. Delta CT (Δ CT) was calculated as the CT of the target minus the CT of β-Actin for each transfection concentration and is plotted against the log of the mass of pDNA transfected (Plotted as mean ± SD).

**Supplemental Figure 2.**
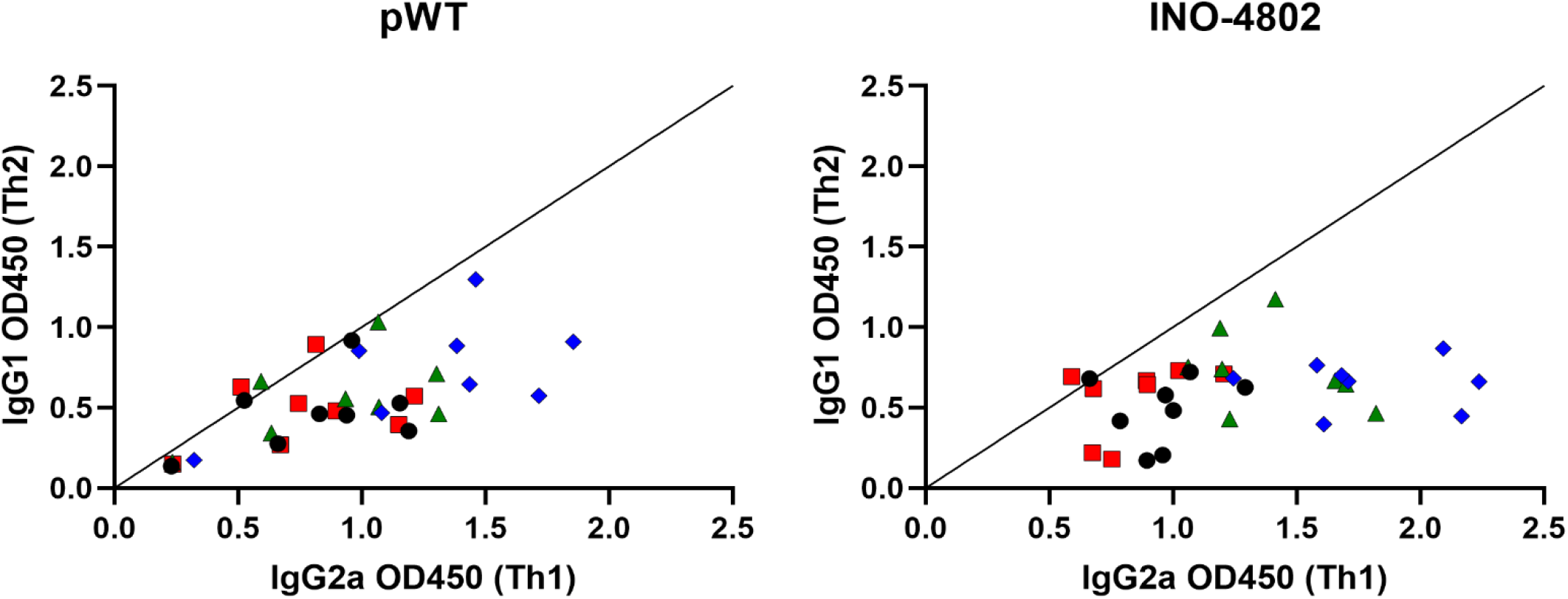
IgG isotype profile of INO-4802 humoral immunogenicity against SARS-CoV-2 VOC. BALB/c mice were immunized on days 0 and 14 with 10 µg of pWT or INO-4802 as described in the methods. Protein antigen binding of IgG2A and IgG1 from mice at day 21 (7 days post second immunization). Data shown represent OD450 nm values for sera at a 1:1350 dilution (linear range of binding for all groups/protein antigens) for each group of mice. Protein antigens are SARS-CoV-2 full length spike proteins (Black circle-WT, Red Square-B.1.1.7, Green Triangle-P.1, Blue Diamond-B.1.351) representing the full mutational profile of each VOC as described in the methods.

**Supplemental Figure 3.**
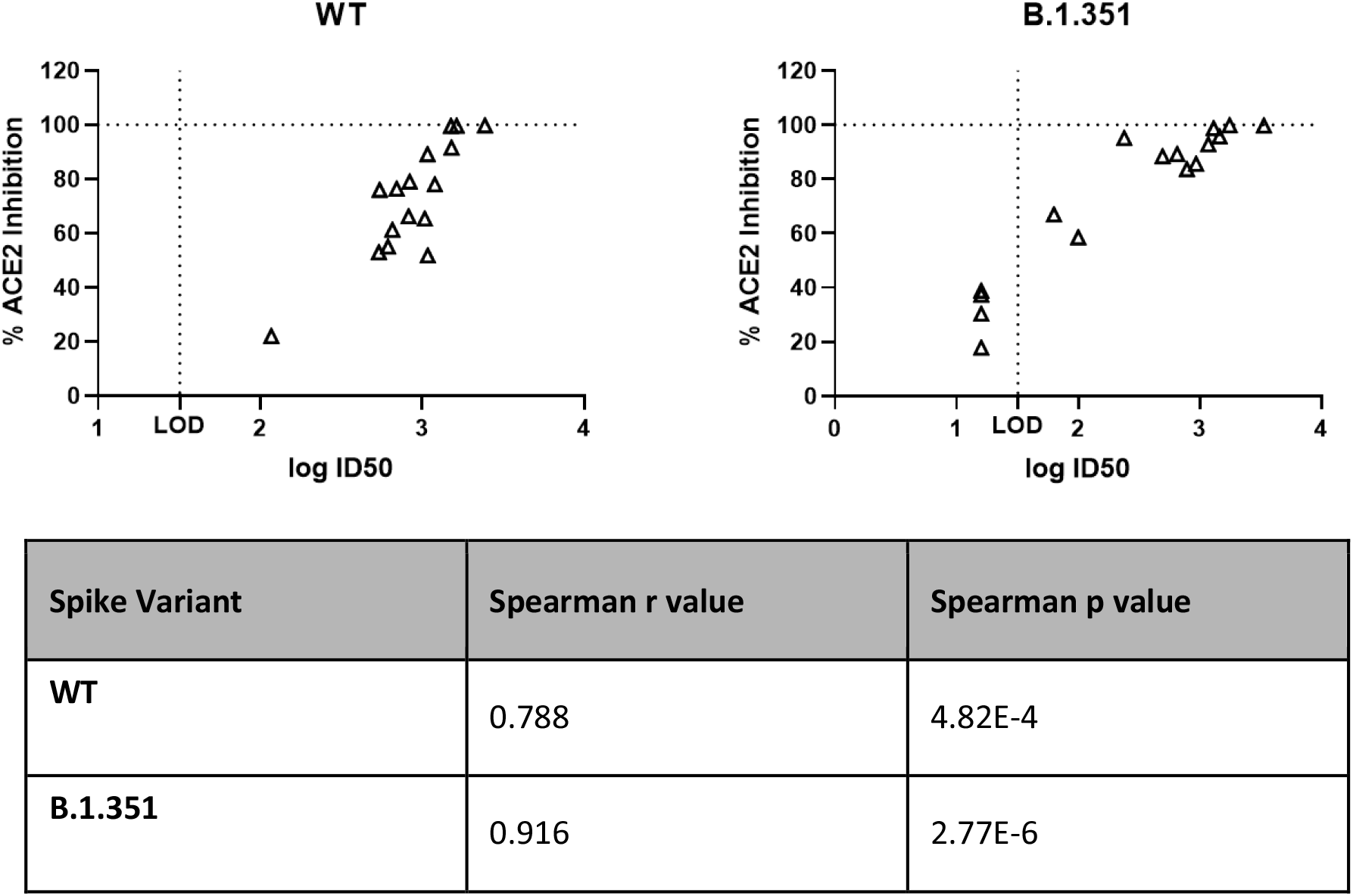
Correlation between pseudovirus and ACE2 blocking assays. Relationship between Pseudoneutralization assay (logID_50_) and percent inhibition of ACE2 binding to SARS-CoV-2 spike S1 protein using day 40 pre-challenge sera samples. Assays represent B.1.351 spike protein and B.1.351 pseudovirus. Correlation analyses were performed using the Spearman method.

**Supplemental Figure 4.**
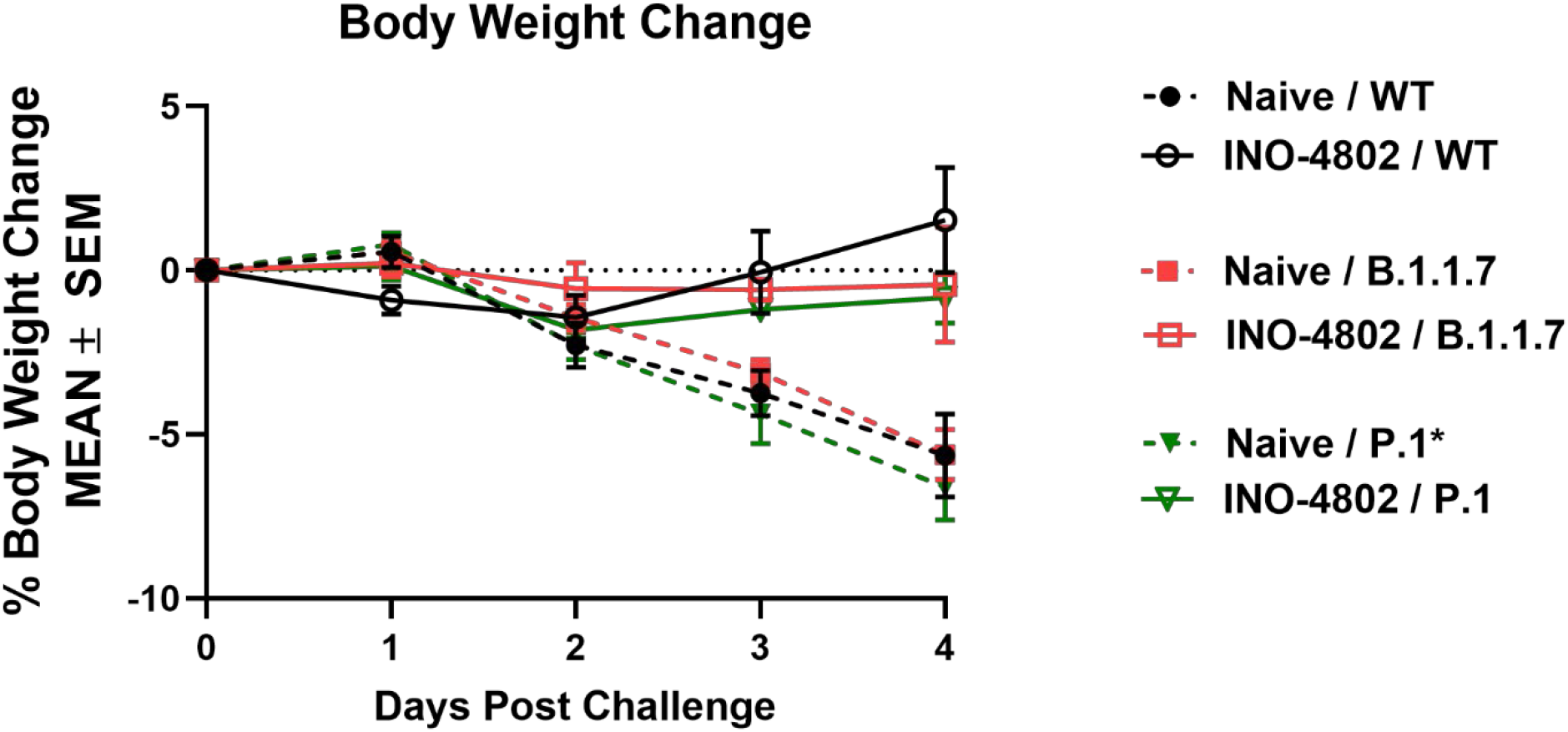
INO-4802 protects Syrian Golden Hamster against challenge with WT and VOCs B.1.1.7, P.1. Animals received ID+EP immunizations with 100 µg INO-4802 on days 0 and 21. Hamsters were challenged on day 55 IN with WT (6000 PFU/hamster USA-WA1/2020), B.1.1.7 (13750 PFU/hamster) or, P.1 (3570 PFU/hamster)* and observed for weight loss. Weight change of INO-4802 vaccinated hamsters compared to unvaccinated animals following challenge with WT, B.1.1.7, P.1. Historical data plotted for P.1 control group

**Supplemental Figure 5.**
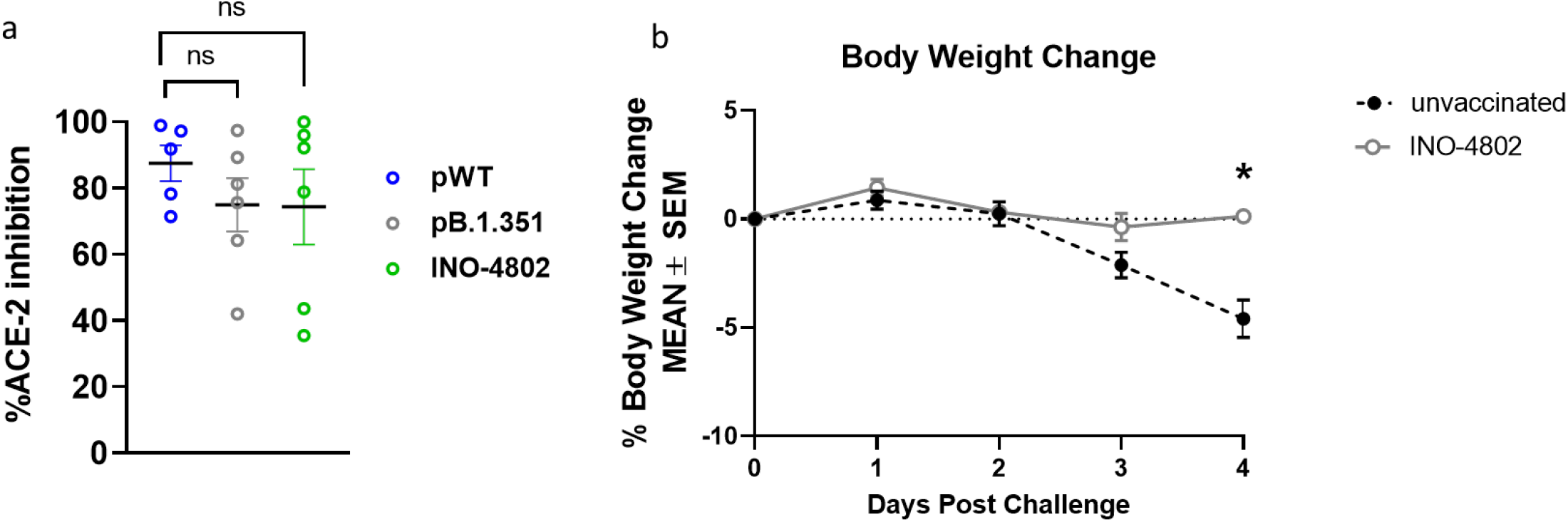
Human ACE2 blocking of B.1.617.2 spike binding by serum from vaccinated hamsters. **a)** Syrian Golden Hamsters received IM+EP immunizations with 10 µg pWT, p.B1.351 or INO-4802 on days 0 and 14. Sera collected on day 22 were tested for capacity to block binding of human ACE-2 to B.1.617.2-spike in an electrochemoluminescent-based ELISA assay (mean %inhibition +/-SEM). Not significant (ns) determined by Welch’s t test. **b)** Animals received ID+EP immunizations with 100 µg INO-4802 on days 0 and 21. Hamsters were challenged on day 70 IN with 6.5×10^3 TCID_50_ B.1.617.2 (obtained from BEI resources: Cat # NR-55612, Lot # 70045240 Titer: 6.5×10^5 TCID_50_/mL (Calu3)) and observed for weight loss. Weight change of INO-4802 vaccinated hamsters compared to unvaccinated animals following challenge with B.1.617.2 (*P < 0.05 determined by Mann-Whitney-test).

## Notes

### Summary of Updates

We provide in depth description of the immunogen design principle and sequence of the construct. We demonstrate the immunogenicity of this construct against a panel of SARS-CoV-2 VOCs in multiple animal models. We demonstrate the efficacy of INO-4802 in a series of hamster challenge studies against multiple SARS-CoV-2 VOCs. We show initial data concerning the neutralizing activity of INO-4800 against the B.1.617.2 (delta) VOC and challenge data.

